# Integrated analysis links *TP53, NOTCH, SLC38A* and 11p with survival in patients with oral tongue squamous cell carcinoma

**DOI:** 10.1101/033829

**Authors:** Neeraja M Krishnan, Saurabh Gupta, Costerwell Khyriem, Vinayak Palve, Amritha Suresh, Gangotri Siddappa, Vikram Kekatpure, Moni Abraham Kuriakose, Binay Panda

**Author notes:** ^*^corresponding author.

## Abstract

Head and neck squamous cell carcinoma (HNSCC) are a heterogeneous group of cancers affecting multiple subsites, including oral cavity. Oral or anterior tongue tumors (OTSCC) are an aggressive group of squamous cell carcinomas, characterized by their early spread to lymph nodes and higher rate of regional failure compared to other oral cavity cancers. There is a rise in the incidence of oral tongue cancer among younger population (<50yrs); many of who lack the typical associated risk factors of alcohol and/or tobacco exposure. In order to carry out an ensemble learning and prediction method with multiple parameters classifying survival, we generated data, on somatic mutations in genes from exome sequencing, immediate upstream and downstream flanking nucleotides of the somatic mutations, DNA methylation, loss of heterozygosity (LOH), copy number variations (CNV), gene expression, significant pathways altered and Human Papilloma Virus (HPV) infection, from 50 OTSCC patients. Results of our analysis identified somatic mutations in *NOTCH2* and/or *TP53*, and/or LOH in 11p to associate with better disease free survival in HPV positive patients (*P* = 0.0254) and not in HPV negative patients (*P* = 0.414). We validated the latter in patients without HPV infection from TCGA cohort (*P* = 0.369, *N =* 17 for TCGA_ OralTongue; *P* = 0.472, *N =* 67 for all TCGA_HNSCC patients). Integrated analysis, including pathways, linked survival with apoptosis and aberrant methylation in *SLC38A8* (*P* = 0.0129).

**Author Summary:** Oral tongue squamous cell carcinomas (OTSCC) are a homogenous group of head and neck tumors characterized with aggressive behavior among younger patients. In this report, we have analysed genetic variants, expression and DNA methylation changes across 50 oral tongue primary tumors along with the Human Papilloma Virus (HPV) infection status in those tumors to identify factors associated with disease free survival. Our data identified somatic mutations in the genes *NOTCH2, TP53* and LOH in 11p, to be significantly associated with better disease free survival in HPV positive patients (*P* = 0.0254), but not in HPV negative patients (*P* = 0.414). We validated the latter using patients without HPV infection from TCGA (*P* = 0.369, *N =* 17 for TCGA_OralTongue; *P* = 0.472, *N =* 67 for all TCGA_HNSCC patients). Integrated analysis linked survival with apoptosis and aberrant methylation in *SLC38A8* (*P* = 0.0129).

## Introduction

Head and neck squamous cell carcinomas (HNSCC) are a heterogeneous group of malignancies with different incidences, mortalities and prognosis for different subsites and are the sixth leading cause of cancer worldwide [1]. In India, they account for almost 30% of all cancer cases [2]. Studies on molecular biology of HNSCC in the past 5yrs have largely been concentrated on cataloging various genetic changes in many cancer types using high-throughput sequencing assays and computational methods [3–7]. Unlike other oral cavity subsites, squamous cell carcinomas of oral/anterior tongue tend to be associated more with younger patients [8, 9], early spread to lymph nodes [8] and a higher regional failure compared to gingivo-buccal cases [10].

Cancer progression and survival are traditionally linked to epidemiological factors, mutations and expression changes in genes, selected primarily on a frequency-based approach that often misses out genuine but low frequency occurrences. Methods based on machine learning have been employed in the past to study effects of genetic variations in diseases [11]. These methods have advantages over parametric methods like logistic regression, which depend on strict and multiple assumptions about the functional form of the model and the effects being modeled, and have limitations in detecting non-linear patterns of interaction [11, 12]. Random forests (RF) by Leo Breiman [13] is a popular machine learning tool for predictive analyses using non-parametric observations, especially when the number of features exceed the number of samples [14]. It has a wide variety of applications in genetic studies [15–17]. However, it is important to estimate errors in such tests, and several methods have been described for estimating errors involving non-parametric observations. The popular ones are *leave-one-out* cross-validation and bootstrapping methods. Bootstrapping methods are generally used to reduce the variability in non-parametric estimators and estimate error rates. The *leave-one-out* cross-validation method has a high variance and estimates errors with an upward bias [18–20]. One of the popular bootstrapping methods, called the .632 method [18], results in a downward bias [19, 20]. This method and other alternatives, thereof, are especially useful for unknown sample sizes and distributions, and are reported to perform better than cross-validation [19–21]. Variable importance is further estimated by permuting values for each feature and computing the average difference in out-of-bag (OOB) error rate before and after the permutation over all trees, normalized by the standard deviation of these differences. Features with larger values for this importance score are ranked as more important than others. The RF algorithm copes well even with highly correlated features and other complex interaction structures, and is able to capture non-linear association patterns between predictors and response [13].

Neighboring nucleotide biases have been reported in previous studies influencing a wide variety of effects including, mutations in mice and human [22], rates and patterns of spontaneous mutations in primates [23], substitutions in human genes [24], and MNU-induced mutations in rice [25]. Neighborhood properties of mutations are also linked to a wide range of biological properties namely, temperature-sensitive mutations [26], regulation of gene expression in human and other primates [27, 28], DNA repair and DNA flanking the region of triplet expansion [29].

Neighborhood properties of mutations was proved to be one of the triggering mechanisms in several human diseases (myotronic distrophy, Huntington's disease, Kennedy's disease and fragile X syndrome [29], and cancer [30]). Hotspots of mutations along with changes in methylation, LOH, CNV and gene expression, present across the genome were discovered in different subsites of head and neck cancer [3–7]. Previous studies link sequence context, such as guanine holes, to pathogenic germline mutations as well as cancer-specific somatic mutations [30]. In fact, specific mutation signatures with the context of certain flanking residues have been identified for some cancers [31, 32]. In the study described here, we used *in house* generated data (OTSCC_MIECL [33] and OTSCC_Methylation (NM Krishnan et al. 2015b *(unpublished))*, on somatic mutations and indels from exome sequencing (*N* = 50) along with data on both 5’ and 3’ flanking nucleotides of the somatic mutations, DNA hypo- and hyper-methylations, LOH, cancer-related pathways specifically altered in tumors along with the data on Human Papilloma Virus (HPV) infection to perform RF individually or in combinations to identify a minimal signature of survival. We used a modified version of .632 bootstrapping method, called .632+ method that corrects the upward bias in the *leave-one-out* and the downward bias in the original .632 bootstrap method [18]. The method used here, the tree learning algorithm, repeatedly bootstraps random subsets of features and samples, and fits a random forest of trees to these, while estimating the average OOB error over the forest. We performed Kaplan-Meier survival probability analyses using predictive parameters from the random forest analyses, and found a combination of molecular markers that are associated with Disease-Free Survival (DFS) among the HPV positive patients, but not in others. We validated these findings using data from The Cancer Genome Atlas (TCGA) project (*N* = 90, oral tongue only, Dataset: TCGA_OralTongue; and *N =* 380 tumors from all subsites of head and neck, Dataset: TCGA_HNSCC).

## Results

We collected tumor and blood/adjacent normal samples from 50 patients diagnosed with OTSCC with informed consent as described previously [33]. In our study, 68% of the patients were alive at the time of analysis and recurrence at primary site was observed in 16% of the patients. Data on survival and recurrence were collected by patient follow up from the date of surgery up to 91 months (the longest duration of survival). In the study group, the median disease-free survival (DFS) was 15 months. HPV subtypes, 16 and 18, infections were detected in 23 out of 50 patients. We divided all the patients into three groups based on the duration of their DFS, group one with DFS less than a year (low, in 38% of patients), group two with DFS between 1–2yrs (med, in 32% of patients) and group three with DFS of >2yrs (high, in 30% of patients). Patients (*N* = 2) were excluded that did not satisfy the study inclusion criteria. Details on the patients with DFS in months, DFS categories, clinical and treatment details and HPV infection, are provided in Supplementary Table 1. We tested the efficiency of predicting survival in the oral tongue squamous cell carcinoma (OTSCC) patients using variable elimination with RF approach, with either one of the following parameters: somatic mutation frequencies within six categories, one and three immediate 5’ and/or 3’ flanking nucleotides of the somatic mutations, somatic mutations in gene bodies, genes affected by somatic mutations, DNA hypo- and hyper-methylations, chromosome arm-level LOH, pathways modified and HPV infection, or in combination (all). For all iterations of all RF analyses, we confirmed that the rankings of variable importances remained highly correlated before and after correcting for multiple hypotheses comparisons using pre- and post-Benjamin-Hochberg false discovery rate (FDR)-corrected *P* values (Supplementary Table 2).

The survival prediction score in our analyses is composed of four components: repC – number of repeatability (confidence) classes, repE – confidence scores of predictors, Sen – sensitivity (number of accurate predictions) and Spec – specificity (classes of accurate predictors). We penalized the score based on repC, and reward for the latter three components.

We found patient-specific survival signatures for various predictor categories (Supplementary Table 3), which included candidate markers known to be associated with survival, like *TP53* and *NOTCH2* genes, and LOH in 11p [34–36]. We conducted a number of single-parameter analyses and a combined-parameter one, which included the top scoring predictors from all the single-parameter analyses.

Non-parametric tests predicted the overall prediction score to be highest for the somatic mutation predictor for all DFS categories (Fig. 1A). In the low DFS category, the 3’ nucleotide flanks ranked second in predicting poor survival with a high score. Overall survival prediction score was also highest with the mutation category predictor for low and high DFS categories, except for the med category, where the somatic mutations ranked first (Supplementary Fig. 1). Following overall analysis, we compared the individual components of the prediction score, and found repE to contribute largely to the overall score for somatic mutation category frequencies (Fig. 1B).

**Fig. 1.**
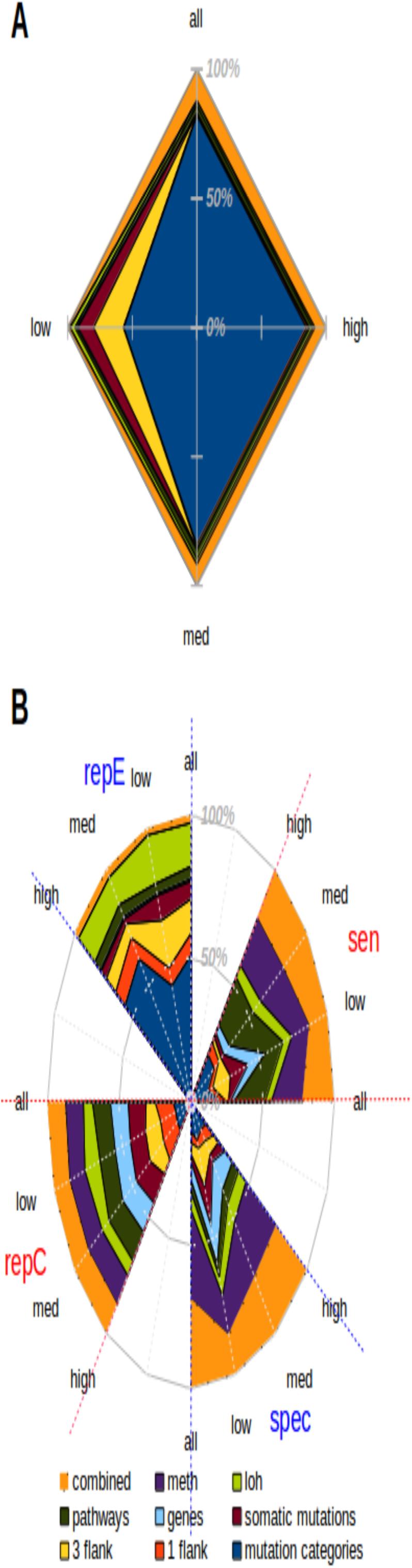
Relative overall and component-wise disease-free survival prediction score plot for single- and combined-parameter analyses. A. Disease-free survival prediction scores are plotted for the all, high (>24 months), med (12–24 months) and low (< 12 months) survival categories as a stacked (%) net chart. The colors represent the single-and combined-parameter from the RF analyses: mutation categories - frequencies of filtered somatic mutations categorized into C->A, C->T, C->G, T->C, T->G & T->A, 1 flank - presence of mutations with a certain 5' and 3' flank combination, 3 flank - presence of mutations with a certain 5' and 3' tri-nucleotide flank combination, somatic mutations - presence of mutations at certain locations coding for genes, genes - genes with mutations, pathways - pathways affected by mutations, loh - chromosomal arms affected by LOH, meth - probes within dmrs, combined - combination of parameters chosen as best predictors in the single-parameter analyses. B. Individual components of disease-free survival prediction scores are plotted for all, high, med and low categories as stacked (%) net chart, for all the single- and combined-parameter analyses. The components are: repC – number of repeatability (confidence) classes, repE – confidence scores of predictors, Sen – sensitivity (number of accurate predictions) and spec – specificity (classes of accurate predictors)

Sensitivity was highest for combined, methylation and pathways, while the former two also had the largest specificity. A similar trend was observed with overall survival as well (Supplementary Fig. 1). We estimated the error in survival prediction using the 0.632+ bootstrapping method [18]. The errors varied across the low, med and high categories of disease-free survival (Supplementary Fig. 2A). The median error was low (<0.3) for the low category for all single-parameter analyses, but was relatively higher (~0.75) for the combined-parameter analyses. The med and high categories had the least median errors (~0.25) for the combined-parameter analyses.

In order to test the specificity of the survival predictors in the OTSCC study, we performed cross-validation using somatic mutation data from three different cancer types (TCGA), either individually or combined. These cancer types were selected based on certain inclusion criteria (see Methods). We found that the survival predictors identified were specific to OTSCC cancer type, against any of the three cancer types, tested individually or pooled (Supplementary Fig. 2B).

For Kaplan-Meier cumulative survival probability analyses on combinations of parameters, we picked *TP53* and *NOTCH2*, and 11p, from gene- and LOH-based single-parameter analysis, respectively. We then assessed survival probability in HPV-background, both in OTSCC and TCGA_HNSCC datasets. However, due to the lack of sufficient number of HPV-positive samples in any of the TCGA_HNSCC subsites, we could only perform survival probability analyses using the OTSCC dataset. The number of samples used for survival probability analyses for various datasets and the exact locations of *TP53* and *NOTCH2* mutations in the internal OTSCC dataset are provided in Supplementary Table 4.

We observed that HPV-positive OTSCC patients, who harbor at least one somatic mutation either in *TP53* or *NOTCH2*, or LOH in 11p, have better disease-free survival (*P* = 0.0254; Fig. 2A). The same is not observed, however, for HPV-negative OTSCC patients (*P* = 0.414; Fig. 2B). We also found that presence of HPV DNA in tumors or somatic mutations/changes in these genes alone does not distinguish disease-free survival (*P* = 0.302; Fig. 2C & *P* = 0.117; Fig. 2D) in OTSCC.

**Fig. 2.**
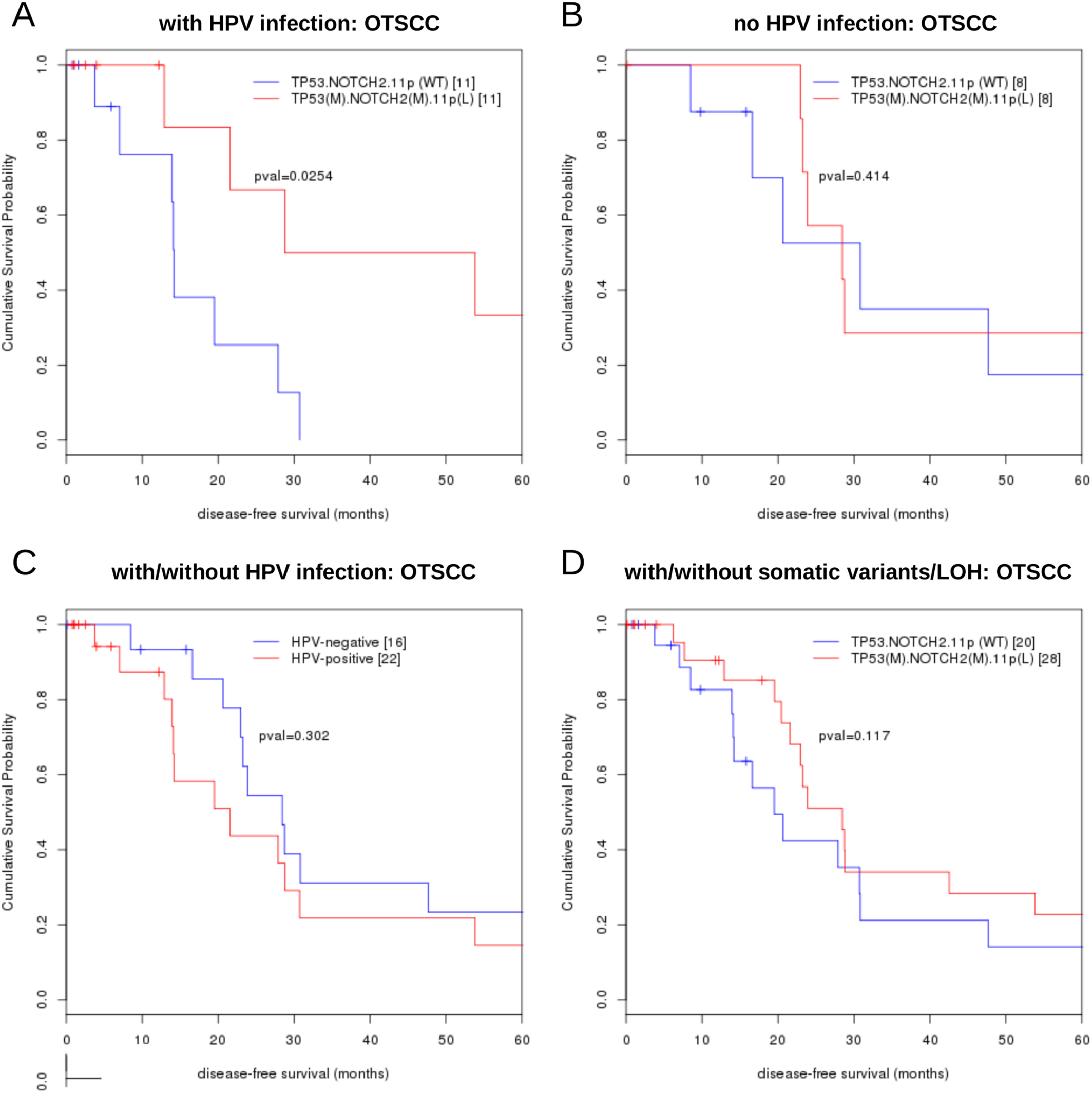
Kaplan-Meier cumulative survival probability curve of combined-parameter analyses. *TP53* and *NOTCH2* variants and 11p LOH combinations against the wild type/non-variant background in patient samples A. with and B. without HPV infection, C. with or without HPV infection, D. *TP53* and *NOTCH2* variants and 11p LOH combination against the wild type/non-variant background. *TP53.NOTCH2.11p* (WT): a combined wild-type background for *TP53, NOTCH2* and 11p; TP53(M).NOTCH2 (M).11p(L): either with *TP53* or *NOTCH2* mutation or mutations in both genes, or 11p LOH or in the presence of all three variants. Significance was assessed using log-rank test.

The TCGA_OralTongue dataset had only one HPV-positive sample (out of 90, where DFS data are available), which harbored a *TP53* mutation but was wild type for *NOTCH2* and 11p, and among all other TCGA_HNSCC HPV-positive samples (*N* = *37)*, 13 patient tumors harbor mutations in *TP53* and/or *NOTCH2* and/or LOH in 11p. However, since these samples were sparsely distributed across multiple sub-sites with different biology and HPV infection rates known to affect survival to varying degrees (for example, a clear link between HPV and survival in oropharyngeal tumors verses no clear data on oral cavity tumors), we did not pool all the HPV-positive patients from the TCGA cohort for analyses. We, however, validated our findings in patients without HPV infection using three other sets of samples from TCGA (*P* = 0.369, 0.723 and 0.472; Fig. 3A-C) with different background of somatic mutations in *TP53* and/or *NOTCH2* and/or LOH in 11p. The first one where we used TCGA samples was the data from the oral tongue subsites only (TCGA_OralTongue, *N =* 17), the second from all head and neck subsites other than in oral tongue (TCGA_HNSCC without OralTongue, *N =* 50) and the third dataset had all subsites of head and neck (TCGA_HNSCC, *N =* 67), where all sample sets were HPV negative. The same combinations of markers were therefore, not significant survival predictors in patients that were HPV negative in all different patient sets. It must be noted that we pooled samples across subsites (TCGA_HNSCC without OralTongue and TCGA_HNSCC), since there was no HPV infection in these cases.

**Fig. 3.**
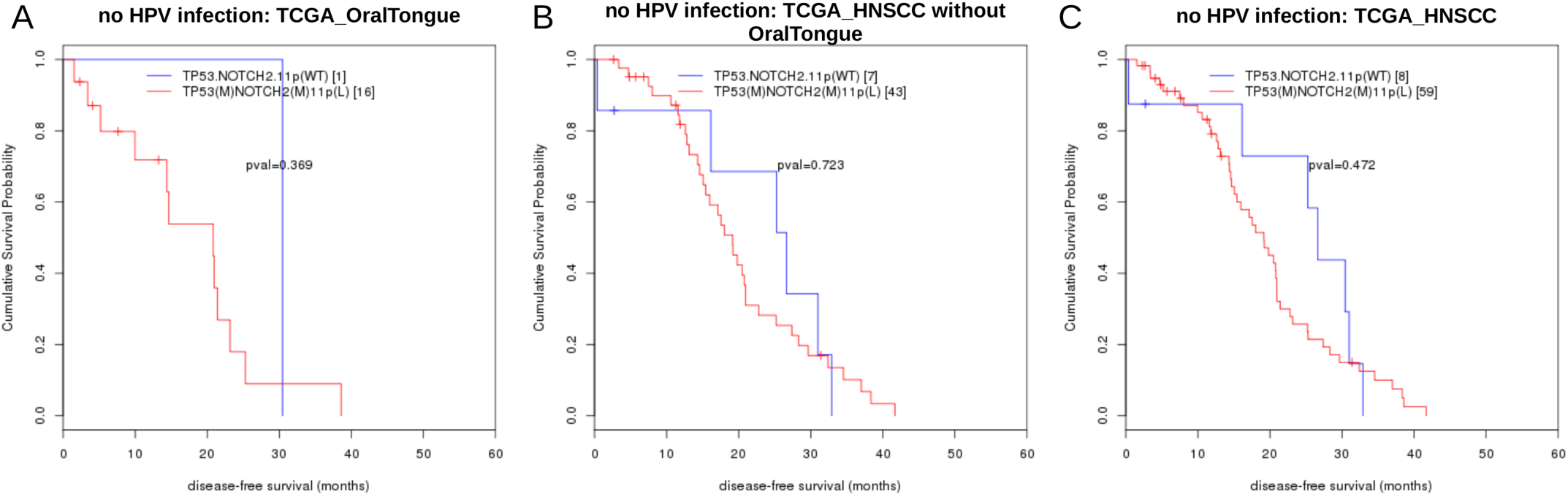
Kaplan-Meier cumulative survival probability curve of combined-parameter analyses predictors. *TP53* and *NOTCH2* variants and 11p LOH combinations against an all three wild type background in patient samples without any HPV infection, in the A. TCGA_OralTongue, B. TCGA_HNSCC without OralTongue and C. TCGA_HNSCC datasets.

In order to determine a minimal predictor set for survival in OTSCC patients, we performed integrated analysis with all predictors in OTSCC patients. We included 120 genes bearing somatic mutations and indels, apoptosis pathway, four methylation probes from whole-genome 450K array (Datasets: OTSCC_MIECL & OTSCC_Methylation), two flanking nucleotides categories, C(T->C)G and T(T->C)G, as top predictors for survival analysis obtained from individual parameter analysis (Supplementary Table 3). For further detailed Kaplan-Meier cumulative survival probability analyses on combinations of parameters, in addition to methylation in *SLC38A8* and nucleotide flanks, we picked *TP53* and *NOTCH2* from the gene-based survival prediction analyses, the former also affecting the apoptosis pathway, along with LOH in the 11p arm, the highest repE-scoring single-parameter analyses and also a known correlate of survival. As shown in Fig. 4, apoptosis pathway, *CpG* island methylation of the *SLC38A8*, somatic mutations in *TP53, NOTCH2* and/or LOH in 11p, forms a minimal predictor for survival in HPV infected OTSCC patients (*P* = 0.0129; Fig. 4A), but not in patients not infected with HPV (*P* = 0.772; Fig. 4B). We could not validate the results from the combined parameter analyses with the TCGA data, as we did not have both the whole-genome 450K DNA methylation data and raw sequence files to extract the flanking nucleotide information for the same tumors.

**Fig. 4.**
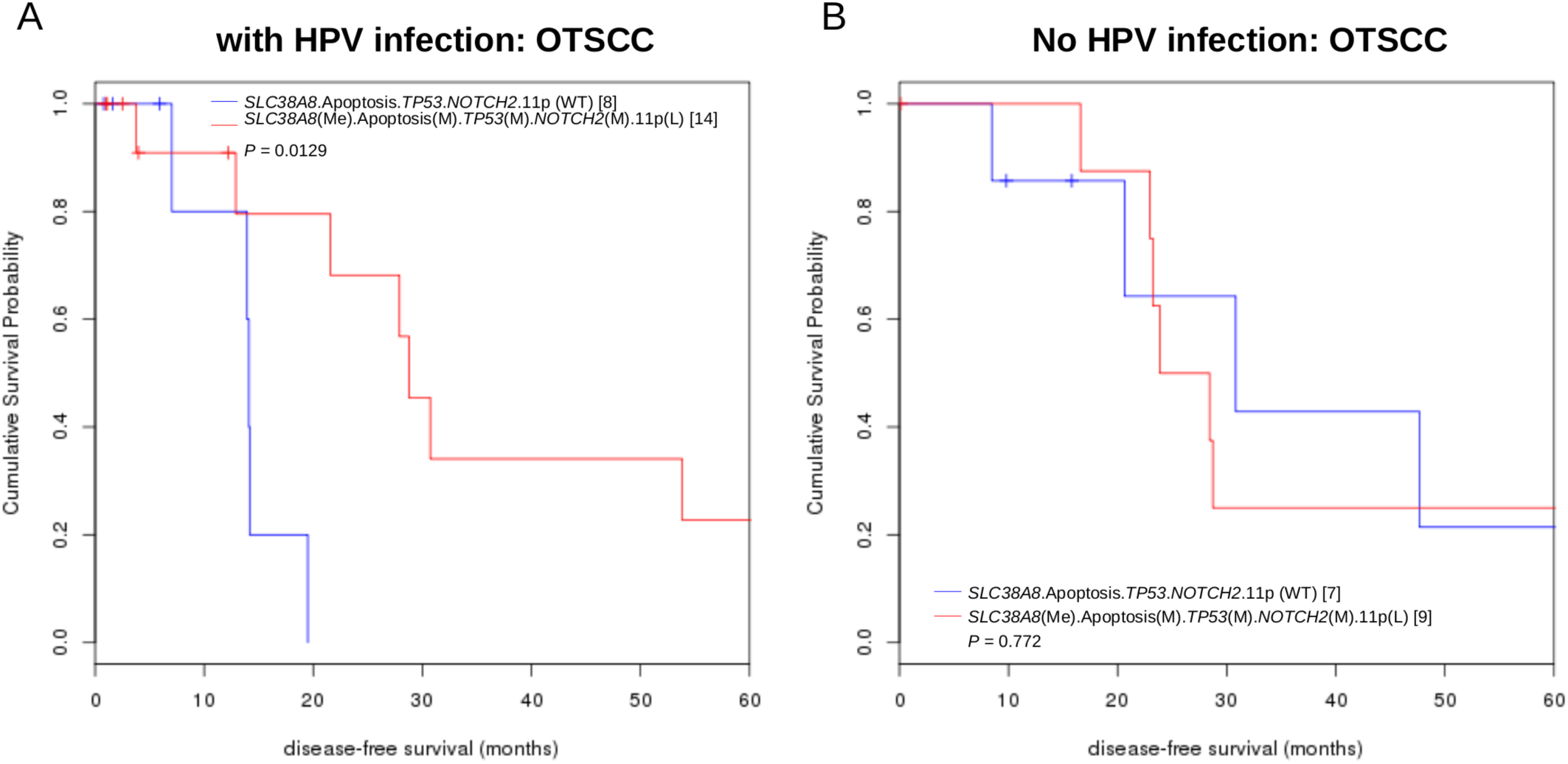
Kaplan-Meier cumulative survival probability curves of individual- and combined-parameter analyses predictors. Significance was assessed using log-rank test. *TP53* and *NOTCH2* variants, 11p LOH, *SLC38A8* methylation and apoptosis pathway variant combinations against an all three wild type control, in OTSCC patients A. with and B. without HPV infection. (SLC38A8.Apoptosis.TP53.NOTCH2.11p)WT: a combined wild-type background for *SLC38A8*, Apoptosis pathway, *TP53, NOTCH2* and 11p; (SLC38A8)Me Apoptosis(M) (TP53)M (NOTCH2)M (11p)L: either one or more of them are perturbed.

Out of the genes and LOH picked from our single parameter RF analysis, we found *TP53, NOTCH2*, and LOH in 11p, along with apoptosis and methylation in *SLC38A8*, as a significant minimal signature of survival in OTSCC patients, where *NOTCH2* mutations were mutually exclusive with *TP53* mutations and/or LOH in 11p.

## Discussion

Earlier survival studies in HNSCC have linked mutations in *TP53* to loss in 3p in HPV negative tumors [37] and identified the role played by mutations in the arachidonic acid metabolism pathway [38]. Currently, the role of genetic and epigenetic changes in the background of HPV infection and their role in survival in HNSCC are lacking. In HNSCC tumors, it has been indicated that HPV plays a limited role outside the oropharynx. In the current study, we investigated the contribution of multiple parameters from genome-wide studies, somatic mutations, indels, immediate flanking nucleotides of somatic mutations, genes and pathways affected by somatic mutations and indels, DNA methylation, LOH, and HPV on disease free survival in patients with oral tongue tumors using a statistical machine learning approach. Synonymous mutations have often, with some evidence, been thought to exert an effect on the function of the gene [39]. For example, a recent report reveals the possibility that many of the synonymous mutations can act as driver mutations in some cancers [40]. We therefore, included them in our analyses.

Neighboring nucleotide biases have been reported in various studies previously [22–25] and were linked to a wide range of properties [26–29]. Sequence context plays an important role in cancer [29–32]. For example, guanine holes are linked to pathogenic cancer-specific mutations [41]. In fact, cancer-type specific mutation signatures with the context of flanking residues have been identified in many cancers [42]. Therefore, we wanted to study the effect of neighboring nucleotides of somatic mutations on survival as an independent predictor, and in combinations with other parameters. In our study, the overlap between annotations from genomic locations of somatic mutations, and the genes and pathways that harbor them provides independent functional validation of the predicted locations and their neighborhood. The contribution of different parameters in three groups of patients was different. In patients with median DFS of <1yr, the contribution of somatic mutations alone was highest but in patients with median DFS >2yrs, it was both the somatic mutations and LOH that contributed near equally (Fig. 1). The overall best prediction of survival suggests that both somatic mutations and LOH are the best predictors (Fig. 2). We found that the flanking nucleotides was a contributing factor and it is possible that the effect of positional context outweighs that of functional context, looking at the lower prediction scores of gene and pathway-based models in comparison to all genomic locations of somatic mutations (133 for genes, 18 for pathways and 3400 for somatic mutations; Supplementary Table 3). The relatively low error for the 'low' survival category for all single-parameter analyses, and for the 'med' and high' categories for all combined-parameter analyses highlights the predictive scope of the two kinds of analyses (Supplementary Figure 2A).

HPV acts through its oncoproteins E6 and E7 that bind and inactivate the cellular tumor suppressor p53 and the retinoblastoma gene product pRb respectively. The degradation of p53 is mediated through ubiquitin pathway. Although HPV has been shown as a good prognostic marker in oropharyngeal cancer, its role in oral cavity tumors is unclear. Past studies have attributed HPV status with therapeutic response and survival in oropharynx tumors [43–45]. Recently, Fakhry *et al*. (2014) showed that patients with oropharyngeal cancer who are p16-positive have significantly improved survival rates when compared with the patients who are p16-negative [46]. Unlike oropharynx, data on the role of HPV in oral cavity carcinomas, including oral tongue, is less conclusive. Previous data showed that the number of patients infected with HPV is much higher in the case of oropharyngeal cancers than those with oral cavity cancers in the United States [47]. HPV prevalence in oral cavity tumors in the United States was shown to be very low (1.5%) [48] to moderate (12.5%) [46]. A recent study in Europe showed 26% of HPV prevalence in oral tongue tumors [45, 46]. Contrary to the data from the United States [48, 49] and closer to the HPV data from Europe [50], studies from India previously reported a high prevalence of HPV in oral cavity tumors, including in oral tongue (anywhere between 18–51% depending on the HPV detection assay used) [51, 52]. Although the reason(s) and importance for such a high prevalence of HPV in our geography is currently not known, the association of HPV with well-differentiated squamous cell carcinoma of oral cavity has been reported in the past [51] in contrast to the oropharyngeal tumors. In a sample size of 50, we found that 38% of the patients were p16 positive and 46% of patients were HPV positive, which is closer to the HPV data obtained earlier in Indian cohorts [51, 52]. A larger sample size and HPV assays with higher sensitivity and specificity in the future will shed more light on the prevalence of HPV in oral cavity tumors. Our results on LOH in 11p and its relationship with survival might link HPV infection in some samples as deletion of 11p was previously shown to enhance viral E6/E7 transcription and virus-mediated cellular transformation in fibroblasts [53]. In our data, we did not find any significant relationship between survival and LOH in 11p when the patients are not stratified based on HPV. It is also known that 11p harbors many putative tumor suppressor genes like *CDKN1C, NUP98, SLC22A18*, and *WT1* [54–58]. Overall, the prognostic markers of survival in OTSCC discovered in our data (in HPV positive and negative background) and validated with the TCGA data (HPV negative background) presents a step forward in understanding the role of HPV in survival of patients with OTSCC.

In our study, we found 19% of the patients that harbor somatic mutations in *TP53* are HPV positive. Even if this was surprising owing to the fact that E6 blocks the function of p53 previous HNSCC studies reported varying numbers of somatic *TP53* mutations in HPV positive background (16% ICGC India [6] and 6.1% TCGA [7]). The reason behind why certain HPV +ve tumors bear mutations in *TP53* is currently unknown. One possibility is that HPV-positive *TP53* mutant tumors may represent a separate group of recurrent/metastatic tumors.

Given the absence of any large study involving patients with oral tongue tumors and the lack of sensitivity of current assays employed to detect HPV, we believe that our data stands unique. As studies involving large number of HPV infected patients in oral tongue are currently missing, we could not validate our findings with a large number of patients. Since we used only patients with confirmed HPV status in all tests involving HPV as a parameter, the effective number of patients was low in all combinations that harbored somatic mutations in *TP53*, and/or, in *NOTCH2* and/or, LOH in 11p. This is one of the drawbacks of our current study. It is interesting to note that presence or absence of HPV infection alone does not associate with better disease free survival (Fig. 2). In the larger cohort of HNSCC patients without HPV infection, *TP53, NOTCH2* and 11p do not play a role in impacting survival in head and neck tumors. These genes link with other important genes and pathways shown to be important in HNSCC and their survival (Supplementary Fig. 3). Our study does not establish the mechanistic insights on why patients with tumors, that are positive in HPV DNA and bear mutations in *TP53* and/or *NOTCH2*, survive longer. This is especially so as it is counter-intuitive to what is known about *TP53* mutations and survival. However, a detailed study on the relationship between mutations in *TP53*, *NOTCH2* and HPV infection in HNSCC is currently lacking, as there are not many tumors described in the literature that bear mutations in *TP53* and/or *NOTCH2* and are positive for HPV DNA. Future studies will shed light on this. Kaplan-Meier analyses using mutation data from other *NOTCH* receptors *(NOTCH1, NOTCH2NL* and *NOTCH3)* did not result in a significant survival signature (Supplementary Table 1), emphasizing the importance of *NOTCH2*. In fact, *NOTCH2* has been shown to act as a driver in HNSCC, previously (Pickering et al. 2014). Like our study, the other sequencing studies also have shown presence of somatic mutations in *NOTCH1* and *NOTCH2* somatic mutations in oral cavity tumors to vary (13% and 5% in [4]; (13% and none in [5]; 16% and none in [3]; 18% and 6% in [6] and 3% and 1% in the oral tongue subsites from a larger TCGA cohort [7]. The frequency of *NOTCH1* and *NOTCH2* non-synonymous mutations in our study is (4% each) is very similar to the oral cavity tumors in some of the above studies. In our study, we used both synonymous and non-synonymous somatic mutations for *NOTCH1* and *NOTCH2*. When we did similar analyses with other *NOTCH* receptors using TCGA data, the role of *NOTCH2* becomes even clear for this cohort (Supplementary Table 1).

## Methods

### Informed consent and Ethics approval

Informed consent was obtained voluntarily from each patient enrolled in the study and ethics approval was obtained from the Institutional Ethics Committees of the Mazumdar Shaw Medical Centre (IRB:NHH/MEC-CL/2014/197) as described elsewhere [33].

### Patient samples used in the study

Details of the blood, matched normal and tumor specimens collected and used in the study are described elsewhere[33]. Only those patients with histologically confirmed squamous cell carcinoma that had at least 70% tumor cells in the specimen were recruited for the study. Fifty treatment-naäve patients who underwent staging according to AJCC criteria, curative intent treatment as per NCCN guidelines involving surgery with or without post-operative adjuvant radiation or chemo-radiation at the Mazumdar Shaw Cancer Centre were accrued for the study. Post-treatment surveillance was carried out by clinical and radiographic examinations as per the NCCN guidelines. DFS was estimated as the maximum follow-up period during which a patient remained free of the disease after treatment.

### Exome sequencing, whole-genome SNP microarray and methylation array

We generated data on somatic mutations and flanking nucleotides from exome sequencing, DNA methylation using Illumina 450K microarrays, LOH and copy number variations using Illumina Omni whole-genome 2.5million SNP microarrays from patients (*N* = 50) with OTSCC to discover signature of survival. Details on cataloguing these variants and the computational pipeline used for discovery of variants are described elsewhere [33]. Somatic variants were narrowed down further to contain only those, which have no read coverage in the matched normal. These somatic variants contained synonymous and non-synonymous mutations, frame-shifted and in frame insertions and deletions. The pre-processed somatic variants for various cancer projects along with the survival information for each patient were used for building models for survival prediction using varSelRF, an R Bioconductor package encoding variable selection from random forests. Two out of fifty patients were not included in further analyses, as they did not pass the study inclusion criteria.

### Feature selection with flanking nucleotides around somatic mutations

We calculated somatic mutation frequencies after classifying them into six mutation categories, and their +/-1 and +/-3 nucleotide flanking sequence neighborhood (Supplementary Table 3). For our study, we have used both the immediate single neighboring nucleotide (+/-1) and three neighboring nucleotides (+/-3) of the somatic mutations. The input dataset for the flank analyses is a set of well-annotated and filtered somatic mutations for OTSCC. We classified mutations into six categories, namely, C->A, C->T, C->G, T->A, T->C and T->G, which would also respectively, encompass the corresponding mutations on the complementary strand, namely, G->T, G->A, G->C, A->T, A->G and A->C. There are 16 possibilities of 1 nucleotide flanking either sides of a somatic mutation, for a given mutation category. For e.g., A.(C->A).A, A.(C->A).C,…,T.(C->A).T. In this manner, we extracted neighboring flanking nucleotide information for the entire OTSCC mutation data and developed models for survival prediction containing 6 categories, and 96 and 24,576 flanks (both at +/-1 and +/-3 flanking nucleotides of the somatic mutations) that define immediate neighborhood of the somatic mutations, on the basis of number of mutations falling within one of the 6 mutation categories, flanked by one and three nucleotides, respectively, on either end. Somatic indels were not included in the flank analyses, due to the variable nature of these events posing a difficulty in determining a contextual neighborhood.

### Other random forest models for feature selection

In addition, we validated the results by building random forest models based on the actual mutations including their genomic location information, their gene and pathway annotations, arm-level LOHs and probe-wise methylation differences. In all thus, we had eight random forest models for variable selection. In the somatic mutations, genes and pathways models, we restricted the somatic mutations to only those that have gene and pathway annotations and performed a set of analyses where we did not restrict it to the gene and pathway information, for comparison. For annotation, mutations were mapped using BEDTools version 2.16 on the ENSEMBL 75 database, for genes, followed by pathway mapping using GraphiteWeb.

### Variable selection

The algorithm performs both backward elimination of variables and selection based on the importance spectrum. About 20% of the least important variables are eliminated iteratively until the current OOB error rate becomes larger than the initial OOB error rate or the OOB error rate in the previous iteration. We performed the above step with 500 iterations, each time with a different seed, and recorded for each, the accuracy of survival prediction with the minimal set of parameters, along with the set size, the actual identity of parameters. We further assessed the number of times a minimal set was repeated over the 500 repetitions. We computed a metric by multiplying the prediction accuracy (sensitivity and specificity) and repeatability, and dividing the product by the number of parameters contained within the predicted set, and reported the set with the highest value for this metric. The Bioconductor package VarSelRF was used for implementation of the variable selection algorithm. The R commands used in our method are provided below.

> *library(randomForest)*
>
> *library(varSelRF)*
>
> *DS=read.table("/home/neeraja/RF/OSCC/TminN.NoFlanks.forRF",header=TRUE,na.string s="NA")*
>
> *DS<-na.omit(DS)*
>
> *for (i in 1:500) {*
>
> *set.seed(i)*
>
> *DS.rf.vsf=varSelRF(DS[, -*
>
> *1],DS[,1],ntree=3000,ntreeIterat=2000,vars.drop.frac=0.2,whole.range =*
>
> *FALSE,keep.forest = TRUE)*
>
> *print(DS.rf.vsf$selected.vars)*
>
> *print(predict(DS.rf.vsf$rf.model,subset(DS[,-1],select=DS.rf.vsf$selected.vars)))*
>
> }

For all iterations of all random forest analyses, all the variable importance were re-computed after correction for multiple comparisons testing using Benjamin-Hochberg test, and it was confirmed that the rankings of variable importance remained highly correlated pre- and post-FDR correction of importance values. The R commands used to re-compute importance values after multiple comparisons testing for the six somatic mutation category RF analyses are provided below.

> *# Selected variables from varSelRF analyses*
>
> *DS.rf.vsf$selected.vars*
>
> *ds.rf <-randomForest(DS[,1] ~., DS, ntree = 3000, norm.votes = FALSE, importance = TRUE)*
>
> *#Importance values before FDR correction*
>
> *ds.rf$importance[ -1,3]*
>
> *#Importance values after FDR correction*
>
> *p.adjust(ds.rf$importance[-1,3], method = "BH", n = 6)*

### .632+ bootstrapping

In order to understand the specificity of the best minimalistic predictors of survival, we estimated the .632+ error rate [18] over 50 bootstrap replicates, for each of our analysis. We used the varSelRFBoot function from the varSelRF Bioconductor package to perform bootstrapping. The .632+ method is described by the following formula:

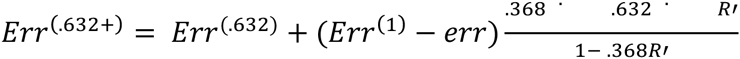

where *Err*^(.632+)^, *Err*^(.632)^, *Err*^(1)^ and *err* are errors estimated by the .632+ method, the original .632 method, *leave-one-out* bootstrap method and *err* represents the error. *R* represents a value between 0 and 1. Another popular error correction method used is *leave-one-out* bootstrap method [21]. The .632+ method was designed to correct the upward bias in the *leave-one-out* and the downward bias in the original .632 bootstrap methods

This method is more accurate compared to the cross validation and the original .632 bootstrapping approaches [18]. It estimates how well the model performs on a subset of the original training set. The .632+ method is an improvement over the original .632 method, where roughly a random sampling of 2/3rds of the data is used as the training set and the rest as the prediction set for each bootstrap replicate, and has a relatively lower upward bias.

### Cross-validation using TCGA data

In order to infer the specificity of survival predictors for the OTSCC study, we performed cross-validation, in addition to the .632+ bootstrapping approach using three different datasets from TCGA (LUSC, GBM and SKCM). The cross-validation was performed while using the three datasets either individually or after combining them. These three cancer types were chosen based on the availability of DFS information, presence of approximately equal number of patients across low (<=12 mo), med (12–24 mo) and high (>=24 mo) DFS categories, and presence of a total of at least 30 patient samples with whole exome sequencing data. We calculated the survival prediction score based on four components: *repC –* number of repeatability (confidence) classes, *repE –* confidence scores of predictors, *Sen –* sensitivity (number of accurate predictions) and *Spec –* specificity (classes of accurate predictors). In order to penalize against too many predictors, we arrived at the following equation:

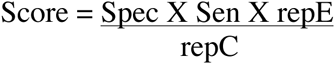

We performed cross-validation for the random forest analyses using six mutation categories and immediate 5′ and 3′ single nucleotide flanks. For these categories, TCGA data possessed the exact same information as the OTSCC data.

### Pathway analysis

Genes, post-annotation of somatic variants, were included to identify signaling pathways and cellular processes through Graphite Web (http://graphiteweb.bio.unipd.it/browse.html) using both KEGG and Reactome databases.

### HPV infection status

We detect HPV positivity using quantitative and droplet digital PCR (ddPCR) assays as described previously[33]. All the samples were processed in triplicates. Genomic DNA from a HPV16 positive (UPCI:SCC-47) and a HPV18 positive (Hep-2) cell line were used as positive and negative controls respectively in these reactions.

### Survival analysis

Survival analysis was carried out using the log-rank test for patients where DFS information was available, and the survival probability distributions for both the groups were estimated using the Kaplan–Meier method and significance assessed using the log-rank test. Significance was always calculated as two-tailed *P* values, on association with DFS of somatic mutations in *TP53* and *NOTCH2*, LOH in 11p, mutations in apoptosis pathway, *CpG* island methylation of *SLC38A8* gene, and HPV in patients as covariates. Observations that pass 95% level of significance are reported. Performing power calculations were not feasible for any of the cohorts, and any efforts to do so resulted in a low power (1-*b*, where *b* is the probability of failing to reject the null hypothesis, given the alternative hypothesis is true).

### Validation with TCGA data

Data with somatic mutations in *TP53, NOTCH2*, LOH in 11p, along with the HPV-infection status and recurrence-free survival were downloaded from the TCGA data portal and the UCSC CancerBrowser (https://genome-cancer.ucsc.edu/proj/site/hgHeatmap/#) for OTSCC (*N* = 90) and for all other subsites of HNSCC (*N* = 290). Using the GISTIC2 threshholded estimates, single-copy deletions in all the 11p associated genes constituted an 11p LOH event. Survival analysis was carried out in various backgrounds, in the presence or absence of mutations (synonymous and non-synonymous) in *TP53* and/or *NOTCH2*, and/or LOH in 11p on both OTSCC and TCGA-_HNSCC data using cumulative probability Kaplan-Meier curves, where censoring of data was performed according to overall survival status, and significance of difference in association determined according to log-rank tests.

### Scripts

All scripts used to deriving data and infer conclusions are provided in Supplementary Scripts.

## SUPPLEMENTARY FILE LEGENDS

The supplementary figures and tables can be downloaded from figshare (http://figshare.com/articles/Survival_Additional_files/1619750).

**Supplementary Table 1:** Calculated *P* values of different combinations of parameters used in Kaplan-Meier analysis and patient details used in the current study. *N1* and *N2* indicate the sample sizes for the two test conditions being compared with respect to differences in association with disease-free survival and *P* (P-value) indicates signficance as assessed by a two-tailed log-rank test. Observations with *N1* <= 5 and/or *N2* <= 5 are excluded from significance assessment. Test conditions with *P* <= 0.05 are indicated in bold. Figure numbers are indicated for the observations used for plotting. Tab 1 contains OTSCC data. Tab 2 contains all TCGA HNSC data. Tab 3 contains patient details used in the OTSCC study.

**Supplementary Table 2:** Correlation between rankings of pre- and post-FDR corrected variable importances. Variable importances were ranked before and after FDR correction using Benjamin-Hochberg test for multiple hypotheses testing. This was done for all random forest iterations yielding the best survival prediction scores.

**Supplementary Table 3:** Patient-wise best scoring predictors of disease-free survival from single- and combined-parameter RF analyses

**Supplementary Table 4:** Details of the various cohorts used in Kaplan-Meier survival probability analyses and the locations and amino acids changes in p53 and Notch2 in OTSCC cohort used in the analyses.

**Supplementary Fig. 1:** Relative A. all and B. component-wise overall survival prediction score plot for single- and combined-parameter analyses. A. Overall survival prediction scores are plotted for the all, high (>24 months), med (12–24 months) and low (< 12 months) categories as a stacked (%) net chart. The colors represent the single-and combined-parameter random forest analyses: mutation categories – frequencies of somatic depth-filtered mutations categorized into C->A, C->T, C->G, T->C, T->G & T->A, 1 flank - presence of mutations with a certain 5' and 3' flank combination, 3 flank – presence of mutations with a certain 5' and 3' tri-nucleotide flank combination, somatic mutations – presence of mutations at certain locations coding for genes, genes – genes with mutations, pathways – pathways affected by mutations, LOH – chromosomal arms affected by LOH, meth – probes within dmrs, combined – combination of parameters chosen as best predictors in the single-parameter analyses. B. Individual components of overall survival prediction scores are plotted for all, high, med and low categories as stacked (%) net chart, for all the single- and combined-parameter analyses. The components are: repC – number of repeatability (confidence) classes, repE – confidence scores of predictors, sen – sensitivity (number of accurate predictions) and spec – specificity (classes of accurate predictors)

**Supplementary Fig. 2:** Performance evaluation of the RF analyses using A. .632+ bootstrapping. .632+ bootstrapping was performed for all iterations of single- and combined-parameter RF runs. The errors were obtained was for all, low, med and high categories of disease-free survival and plotted as box-and-whisker plots using R ggplot *(see* Supplementary Scripts). B. Performance evaluation of the RF analyses using cross-validation. Cross validation was performed using the OTSCC mutation categories (1) and 1 flank (2) data as training sets and similar data from three cancer types, individually, namely, LUSC, GBM and SKCM from TCGA, or all combined, as the prediction sets. Individual components of DFS prediction scores are plotted for all, high, med and low categories as stacked (%) net chart. The components are: repC – number of repeatability (confidence) classes, repE – confidence scores of predictors, sen – sensitivity (number of accurate predictions) and spec – specificity (classes of accurate predictors).

**Supplementary Fig. 3:** Pathway mapping with interactions of key predictors of disease free survival in OTSCC.

Supplementary Scripts: Scripts used to deriving data and infer conclusions.

## REFERENCES

1. Ferlay J, Shin HR, Bray F, Forman D, Mathers C, Parkin DM. Estimates of worldwide burden of cancer in 2008: GLOBOCAN 2008. International journal of cancer Journal international du cancer. 2010;127(12):2893–917. doi: 10.1002/ijc.25516. PubMed PMID: 21351269.

2. Mishra A, Meherotra R. Head and neck cancer: global burden and regional trends in India. Asian Pacific journal of cancer prevention: APJCP. 2014;15(2):537–50. PubMed PMID: 24568456.

3. Agrawal N, Frederick MJ, Pickering CR, Bettegowda C, Chang K, Li RJ, et al. Exome sequencing of head and neck squamous cell carcinoma reveals inactivating mutations in NOTCH1. Science. 2011;333(6046): 1154–7. Epub 2011/07/30. doi: 10.1126/science.1206923. PubMed PMID: 21798897; PubMed Central PMCID: PMC3162986.

4. Stransky N, Egloff AM, Tward AD, Kostic AD, Cibulskis K, Sivachenko A, et al. The mutational landscape of head and neck squamous cell carcinoma. Science. 2011;333(6046):1157–60. doi: 10.1126/science.1208130. PubMed PMID: 21798893; PubMed Central PMCID: PMC3415217.

5. Pickering CR, Zhang J, Yoo SY, Bengtsson L, Moorthy S, Neskey DM, et al. Integrative genomic characterization of oral squamous cell carcinoma identifies frequent somatic drivers. Cancer discovery. 2013;3(7):770–81. doi: 10.1158/2159-8290.CD-12-0537. PubMed PMID: 23619168; PubMed Central PMCID: PMC3858325.

6. India Project Team of the International Cancer Genome C. Mutational landscape of gingivo-buccal oral squamous cell carcinoma reveals new recurrently-mutated genes and molecular subgroups. Nature communications. 2013;4:2873. doi: 10.1038/ncomms3873. PubMed PMID: 24292195; PubMed Central PMCID: PMC3863896.

7. Cancer Genome Atlas N. Comprehensive genomic characterization of head and neck squamous cell carcinomas. Nature. 2015;517(7536):576–82. doi: 10.1038/nature14129. PubMed PMID: 25631445; PubMed Central PMCID: PMC4311405.

8. Llewellyn CD, Johnson NW, Warnakulasuriya KA. Risk factors for squamous cell carcinoma of the oral cavity in young people--a comprehensive literature review. Oral oncology. 2001;37(5):401–18. PubMed PMID: 11377229.

9. Kuriakose M, Sankaranarayanan M, Nair MK, Cherian T, Sugar AW, Scully C, et al. Comparison of oral squamous cell carcinoma in younger and older patients in India. European journal of cancer Part B, Oral oncology. 1992;28B(2): 113–20. PubMed PMID: 1306728.

10. Pathak KA, Das AK, Agarwal R, Talole S, Deshpande MS, Chaturvedi P, et al. Selective neck dissection (I-III) for node negative and node positive necks. Oral oncology. 2006;42(8):837–41. doi: 10.1016/j.oraloncology.2005.12.002. PubMed PMID: 16730221.

11. Moore JH, Asselbergs FW, Williams SM. Bioinformatics challenges for genome-wide association studies. Bioinformatics. 2010;26(4):445–55. doi: 10.1093/bioinformatics/btp713. PubMed PMID: 20053841; PubMed Central PMCID: PMC2820680.

12. McKinney BA, Reif DM, Ritchie MD, Moore JH. Machine learning for detecting gene-gene interactions: a review. Applied bioinformatics. 2006;5(2):77–88. PubMed PMID: 16722772; PubMed Central PMCID: PMC3244050.

13. Breiman L. Random Forests. Schapire RE, editor. The Netherlands: Kliwer Academic Publishers; 2001. 28 p.

14. Jia P, Wang Q, Chen Q, Hutchinson KE, Pao W, Zhao Z. MSEA: detection and quantification of mutation hotspots through mutation set enrichment analysis. Genome biology. 2014;15(10):489. doi: 10.1186/PREACCEPT-1822061407140231. PubMed PMID: 25348067; PubMed Central PMCID: PMC4226881.

15. Jiang R, Tang W, Wu X, Fu W. A random forest approach to the detection of epistatic interactions in case-control studies. BMC bioinformatics. 2009;10 Suppl 1:S65. doi: 10.1186/1471-2105-10-S1-S65. PubMed PMID: 19208169; PubMed Central PMCID: PMC2648748.

16. Lunetta KL, Hayward LB, Segal J, Van Eerdewegh P. Screening large-scale association study data: exploiting interactions using random forests. BMC genetics. 2004;5:32. doi: 10.1186/1471-2156-5-32. PubMed PMID: 15588316; PubMed Central PMCID: PMC545646.

17. Pan Q, Hu T, Malley JD, Andrew AS, Karagas MR, Moore JH. A system-level pathway-phenotype association analysis using synthetic feature random forest. Genetic epidemiology. 2014;38(3):209–19. doi: 10.1002/gepi.21794. PubMed PMID: 24535726.

18. Tibshirani R, Efron B. Improvements on Cross-Validation: The. 632 + Bootstrap Method. Journal of the American Statistical Association. 1997;92(438): 13.

19. Boulesteix A-L, Janitza S, Kruppa J, König IR. Overview of random forest methodology and practical guidance with emphasis on computational biology and bioinformatics. WIREs Data Mining Knowl Discov. 2012;2:15.

20. Boulesteix AL, Janitza S, Hapfelmeier A, Van Steen K, Strobl C. Letter to the Editor: On the term 'interaction' and related phrases in the literature on Random Forests. Briefings in bioinformatics. 2015;16(2):338–45. doi: 10.1093/bib/bbu012. PubMed PMID: 24723569; PubMed Central PMCID: PMCPMC4364067.

21. Efron B. Estimating the Error Rate of a Prediction Rule. Journal of the American Statistical Association. 1983;78:16.

22. Zhang F, Zhao Z. The influence of neighboring-nucleotide composition on single nucleotide polymorphisms (SNPs) in the mouse genome and its comparison with human SNPs. Genomics. 2004;84(5):785–95. doi: 10.1016/j.ygeno.2004.06.015. PubMed PMID: 15475257.

23. Blake RD, Hess ST, Nicholson-Tuell J. The influence of nearest neighbors on the rate and pattern of spontaneous point mutations. Journal of molecular evolution. 1992;34(3):189–200. PubMed PMID: 1588594.

24. Krawczak M, Ball EV, Cooper DN. Neighboring-nucleotide effects on the rates of germ-line single-base-pair substitution in human genes. American journal of human genetics. 1998;63(2):474–88. doi: 10.1086/301965. PubMed PMID: 9683596; PubMed Central PMCID: PMC1377306.

25. Suzuki T, Moriguchi K, Tsuda K, Eiguchi M, Kumamaru T, Satoh H, et al. Neighboring nucleotide bias around MNU-induced mutations in rice.. Rice Genetics NewsLetter. 2008;25:90.

26. Lockwood S, Krishnamoorthy B, Ye P. Neighborhood properties are important determinants of temperature sensitive mutations. PloS one. 2011;6(12):e28507. doi: 10.1371/journal.pone.0028507. PubMed PMID: 22164302; PubMed Central PMCID: PMC3229608.

27. De S, Babu MM. Genomic neighbourhood and the regulation of gene expression. Current opinion in cell biology. 2010;22(3):326–33. doi: 10.1016/j.ceb.2010.04.004. PubMed PMID: 20493676.

28. De S, Teichmann SA, Babu MM. The impact of genomic neighborhood on the evolution of human and chimpanzee transcriptome. Genome research. 2009;19(5):785–94. doi: 10.1101/gr.086165.108. PubMed PMID: 19233772; PubMed Central PMCID: PMC2675967.

29. Polak P, Lawrence MS, Haugen E, Stoletzki N, Stojanov P, Thurman RE, et al. Reduced local mutation density in regulatory DNA of cancer genomes is linked to DNA repair. Nature biotechnology. 2014;32(1):71–5. Epub 2013/12/18. doi: 10.1038/nbt.2778. PubMed PMID: 24336318.

30. Jia P, Pao W, Zhao Z. Patterns and processes of somatic mutations in nine major cancers. BMC medical genomics. 2014;7:11. doi: 10.1186/1755-8794-7-11. PubMed PMID: 24552141; PubMed Central PMCID: PMC3942057.

31. Nik-Zainal S, Alexandrov LB, Wedge DC, Van Loo P, Greenman CD, Raine K, et al. Mutational processes molding the genomes of 21 breast cancers. Cell. 2012;149(5):979–93. Epub 2012/05/23. doi: 10.1016/j.cell.2012.04.024. PubMed PMID: 22608084; PubMed Central PMCID: PMC3414841.

32. Taylor BJ, Nik-Zainal S, Wu YL, Stebbings LA, Raine K, Campbell PJ, et al. DNA deaminases induce break-associated mutation showers with implication of APOBEC3B and 3A in breast cancer kataegis. eLife. 2013;2:e00534. Epub 2013/04/20. doi: 10.7554/eLife.00534. PubMed PMID: 23599896; PubMed Central PMCID: PMC3628087.

33. Krishnan NM, Gupta S, Palve V, Varghese L, Pattnaik S, Jain P, et al. Integrated analysis of oral tongue squamous cell carcinoma identifies key variants and pathways linked to risk habits, HPV, clinical parameters and tumor recurrence [version 1; referees: 1 approved] F1000Research 2015;4:1215. doi: doi: 10.12688/f1000research.7302.1.

34. Poeta ML, Manola J, Goldwasser MA, Forastiere A, Benoit N, Califano JA, et al. TP53 mutations and survival in squamous-cell carcinoma of the head and neck. The New England journal of medicine. 2007;357(25):2552–61. doi: 10.1056/NEJMoa073770. PubMed PMID: 18094376; PubMed Central PMCID: PMCPMC2263014.

35. Xu J, Song F, Jin T, Qin J, Wu J, Wang M, et al. Prognostic values of Notch receptors in breast cancer. Tumour biology: the journal of the International Society for Oncodevelopmental Biology and Medicine. 2015. doi: 10.1007/s13277-015-3961-6. PubMed PMID: 26323259.

36. Attiyeh EF, London WB, Mosse YP, Wang Q, Winter C, Khazi D, et al. Chromosome 1p and 11q deletions and outcome in neuroblastoma. The New England journal of medicine. 2005;353(21):2243–53. Epub 2005/11/25. doi: 10.1056/NEJMoa052399. PubMed PMID: 16306521.

37. Gross AM, Orosco RK, Shen JP, Egloff AM, Carter H, Hofree M, et al. Multi-tiered genomic analysis of head and neck cancer ties TP53 mutation to 3p loss. Nature genetics. 2014;46(9):939–43. doi: 10.1038/ng.3051. PubMed PMID: 25086664; PubMed Central PMCID: PMC4146706.

38. Biswas NK, Das S, Maitra A, Sarin R, Majumder PP. Somatic mutations in arachidonic acid metabolism pathway genes enhance oral cancer post-treatment disease-free survival. Nature communications. 2014;5:5835. doi: 10.1038/ncomms6835. PubMed PMID: 25517499.

39. Supek F. The Code of Silence: Widespread Associations Between Synonymous Codon Biases and Gene Function. Journal of molecular evolution. 2015. doi: 10.1007/s00239-015-9714-8. PubMed PMID: 26538122.

40. Supek F, Minana B, Valcarcel J, Gabaldon T, Lehner B. Synonymous mutations frequently act as driver mutations in human cancers. Cell. 2014;156(6):1324–35. doi: 10.1016/j.cell.2014.01.051. PubMed PMID: 24630730.

41. Bacolla A, Temiz NA, Yi M, Ivanic J, Cer RZ, Donohue DE, et al. Guanine holes are prominent targets for mutation in cancer and inherited disease. PLoS genetics. 2013;9(9):e1003816. doi: 10.1371/journal.pgen.1003816. PubMed PMID: 24086153; PubMed Central PMCID: PMC3784513.

42. Roberts SA, Gordenin DA. Hypermutation in human cancer genomes: footprints and mechanisms. Nature reviews Cancer. 2014;14(12):786–800. PubMed PMID: 25568919; PubMed Central PMCID: PMC4280484.

43. Fakhry C, Westra WH, Li S, Cmelak A, Ridge JA, Pinto H, et al. Improved survival of patients with human papillomavirus-positive head and neck squamous cell carcinoma in a prospective clinical trial. Journal of the National Cancer Institute. 2008;100(4):261–9. doi: 10.1093/jnci/djn011. PubMed PMID: 18270337.

44. Jo S, Juhasz A, Zhang K, Ruel C, Loera S, Wilczynski SP, et al. Human papillomavirus infection as a prognostic factor in oropharyngeal squamous cell carcinomas treated in a prospective phase II clinical trial. Anticancer research. 2009;29(5):1467–74. PubMed PMID: 19443352; PubMed Central PMCID: PMC3582681.

45. Wilson DD, Crandley EF, Sim A, Stelow EB, Majithia N, Shonka DC, Jr., et al. Prognostic significance of p16 and its relationship with human papillomavirus in pharyngeal squamous cell carcinomas. JAMA otolaryngology--head & neck surgery. 2014;140(7):647–53. doi: 10.1001/jamaoto.2014.821. PubMed PMID: 24876098.

46. Fakhry C, Zhang Q, Nguyen-Tan PF, Rosenthal D, El-Naggar A, Garden AS, et al. Human papillomavirus and overall survival after progression of oropharyngeal squamous cell carcinoma. Journal of clinical oncology: official journal of the American Society of Clinical Oncology. 2014;32(30):3365–73. doi: 10.1200/JCO.2014.55.1937. PubMed PMID: 24958820; PubMed Central PMCID: PMC4195851.

47. Walline HM, Komarck C, McHugh JB, Byrd SA, Spector ME, Hauff SJ, et al. High-risk human papillomavirus detection in oropharyngeal, nasopharyngeal, and oral cavity cancers: comparison of multiple methods. JAMA otolaryngology--head & neck surgery. 2013;139(12): 1320–7. doi: 10.1001/jamaoto.2013.5460. PubMed PMID: 24177760; PubMed Central PMCID: PMC4049419.

48. Lingen MW, Xiao W, Schmitt A, Jiang B, Pickard R, Kreinbrink P, et al. Low etiologic fraction for high-risk human papillomavirus in oral cavity squamous cell carcinomas. Oral oncology. 2013;49(1): 1 -8. doi: 10.1016/j.oraloncology.2012.07.002. PubMed PMID: 22841678.

49. Liang XH, Lewis J, Foote R, Smith D, Kademani D. Prevalence and significance of human papillomavirus in oral tongue cancer: the Mayo Clinic experience. Journal of oral and maxillofacial surgery: official journal of the American Association of Oral and Maxillofacial Surgeons. 2008;66(9):1875–80. doi: 10.1016/j.joms.2008.04.009. PubMed PMID: 18718395.

50. Garcia-de Marcos JA, Perez-Zafrilla B, Arriaga A, Arroyo-Rodriguez S, Poblet E. Human papillomavirus in carcinomas of the tongue: clinical and prognostic implications. International journal of oral and maxillofacial surgery. 2014;43(3):274–80. doi: 10.1016/j.ijom.2013.10.016. PubMed PMID: 24268899.

51. Elango KJ, Suresh A, Erode EM, Subhadradevi L, Ravindran HK, Iyer SK, et al. Role of human papilloma virus in oral tongue squamous cell carcinoma. Asian Pacific journal of cancer prevention: APJCP. 2011;12(4):889–96. PubMed PMID: 21790221.

52. Ramshankar V, Soundara VT, Shyamsundar V, Ramani P, Krishnamurthy A. Risk stratification of early stage oral tongue cancers based on HPV status and p16 immunoexpression. Asian Pacific journal of cancer prevention: APJCP. 2014;15(19):8351–9. PubMed PMID: 25339028.

53. Smits PH, Smits HL, Jebbink MF, ter Schegget J. The short arm of chromosome 11 likely is involved in the regulation of the human papillomavirus type 16 early enhancer-promoter and in the suppression of the transforming activity of the viral DNA. Virology. 1990;176(1):158–65. PubMed PMID: 2158686.

54. Chu SH, Feng DF, Ma YB, Zhang H, Zhu ZA, Li ZQ, et al. Promoter methylation and downregulation of SLC22A18 are associated with the development and progression of human glioma. Journal of translational medicine. 2011;9:156. doi: 10.1186/1479-5876-9-156. PubMed PMID: 21936894; PubMed Central PMCID: PMC3184631.

55. Borriello A, Caldarelli I, Bencivenga D, Criscuolo M, Cucciolla V, Tramontano A, et al. p57(Kip2) and cancer: time for a critical appraisal. Molecular cancer research: MCR. 2011;9(10): 1269–84. doi: 10.1158/1541-7786.MCR-11-0220. PubMed PMID: 21816904.

56. Singer S, Zhao R, Barsotti AM, Ouwehand A, Fazollahi M, Coutavas E, et al. Nuclear pore component Nup98 is a potential tumor suppressor and regulates posttranscriptional expression of select p53 target genes. Molecular cell. 2012;48(5):799–810. doi: 10.1016/j.molcel.2012.09.020. PubMed PMID: 23102701; PubMed Central PMCID: PMC3525737.

57. Huff V. Wilms’ tumours: about tumour suppressor genes, an oncogene and a chameleon gene. Nature reviews Cancer. 2011;11(2): 111–21. doi: 10.1038/nrc3002. PubMed PMID: 21248786.

58. Hohenstein P, Hastie ND. The many facets of the Wilms' tumour gene, WT1. Human molecular genetics. 2006;15 Spec No 2:R196–201. doi: 10.1093/hmg/ddl196. PubMed PMID: 16987884.

